# Multiple nuclei means multiple chromosome sets in *Botrytis cinerea* and *Neurospora crassa*

**DOI:** 10.64898/2026.03.14.711691

**Authors:** Dian Zheng, Jan A.L. van Kan, Ben Auxier

**Affiliations:** Laboratory of Phytopathology, Wageningen University & Research; Laboratory of Genetics, Wageningen University & Research

## Abstract

We often think of fungi as mysterious organisms that do not follow the general principles of other eukaryotes. Thus, when exciting results are found, these organisms do not always receive the rigorous level of scrutiny seen in other fields. For many fungal species, dispersal and reproduction relies on spores, either sexual or asexual. These spores can either have a single nucleus, or multiple nuclei, and the purpose of these presumably mitotic copies was unclear. Recently it was described that the multiple nuclei in these spores are not mitotic duplicates, but instead they share a single haploid set of chromosomes distributed across nuclei. Here, we provide fluorescent microscopy and UV mutagenesis data that is inconsistent with this hypothesis. Contrasting these previous results, we observe multiple sets of chromosomes in spores of both *B. cinerea* and *N. crassa*. We also observed a strong linear relationship between the number of nuclei in spores and the total acriflavine fluorescence, further supporting mitotic copies. Genome sequencing of colonies arising from UV-irradiated colonies also recovered variants at intermediate frequences, instead of the fixed 100% expected from the new model proposed. This evidence suggests that, as long suspected, these nuclei are indeed mitotic copies, and that a re-evaluation of fungal biology is not currently necessary.

## Introduction

A common observation in fungi is that the dispersal propagules, spores, contain multiple nuclei (Horton, 2006; Yuill, 1950), ranging into the thousands in the case of arbuscular mycorrhizal fungi (Roper et al., 2011). In some cases, like some mushrooms and plant pathogenic rust fungi, having multiple nuclei allows two genotypes to persist in a dikaryotic state and disperse together to ensure reproductive success [cite]. In some cases, the fungal individual can be composed of multiple genotypes, called a heterokaryon, and the spores produced can have nuclei representing the different genotypes present in the colony that produced the spores. However, outside of these heterokaryotic individuals and dikaryotic species the nuclei in multinucleate spores have been assumed to arise from mitotic divisions. Due to this mitotic, and thus clonal, relationship it has been a universal assumption that across these different nuclear organizations each nucleus contains a full complement of all essential chromosomes.

The behavior of nuclei within multinucleate spores has been best studied in the fungus *Neurospora crassa*. This fungus was extensively used by Beadle and Tatum to generate mutants when demonstrating the “one gene – one enzyme” principle (Beadle & Tatum, 1941). In this species it was recognized from very early observations that these multinucleate conidia can carry different alleles of a given gene, and thus must have multiple copies of a given chromosome, a condition called heterokaryosis (Dodge, 1928). These nuclei seem to be combined in a random fashion, although some biases may exist (Atwood & Mukai, 1955), meaning that even if the underlying colony is a heterokaryon some spores will be homokaryotic, sharing all alleles, because the of random nuclear segregation (Klein, 1958; Prout et al., 1953). In fact, these heterokaryotic spores were a major problem for Beadle and Tatum, as mutagenized spores frequently carried both wild-type and mutant alleles for a particular metabolic gene, that they developed a tedious sexual crossing method to purify mutants (Beadle & Tatum, 1945). More currently, the fact that these different nuclei contain different alleles of the same gene is readily observed both through evolutionary experiments where a sup-population of cheaters persists through this heterokaryotic state (Bastiaans et al., 2016), as well as direct visualization of nuclei that encode different fluorescent proteins at the homologous insertion point (Mela & Glass, 2023).

Recently, this concept has been challenged, instead suggesting these nuclei actually together have a single chromosome set and are effectively haploid. With evidence from the genetic model *N. crassa* and the plant pathogenic fungi *Sclerotinia sclerotiorum* and *Botrytis cinerea* various lines of evidence are shown that chromosomes are distributed over multiple nuclei. Using a mix of microscopy, flow cytometry, and genetic information, they suggest that through undescribed mechanisms, these nuclei are coordinated such that the chromosomal complement remains constant at 1N. Not only in the spores, it is also suggested that in the mycelia these chromosomes are still distributed between nuclei, although meiosis proceeds normally (Tian et al., 2025).

The new biology suggested by this new model would overturn much of what is known about fungal biology. Helpfully, this model also makes several strong predictions. Here, we test these predictions. Contrasting this new model, we find a strong linear relationship between the number of nuclei in a spore and the total DNA content across both *B. cinerea* and *N. crassa*, indicating that nuclei are indeed mitotic copies. Observations of prepared protoplasts also reveal multiple chromosome sets present in a single spore, likely related to preparation methodology. Testing the effect of a mutagen, ultraviolet light, we failed to recover the homokaryotic mutants predicted by this model. Our combined microscopic and genetic data suggest that these previous observations are artifactual and that, in multinucleate spores, each nucleus contains a complete chromosomal complement as previously assumed.

## Results

### Microscopy of chromosomes released from *Botrytis cinerea* conidial germlings

A key prediction of this model is that between the multiple nuclei of a spore, only a single haploid set will be present. To observe individual chromosomes, we used a method similar to the protocol described for *Sclerotinia* sexual spores (Xu et al., 2025). By germinating spores in the presence of nocodazole, mitosis is blocked but the germination process allows the cell wall to be enzymatically digested, producing protoplasts. To improve efficiency, we extended the germination time (in continuous presence of nocodazole), which yielded sufficient protoplasts for subsequent analysis. When protoplast suspensions were dropped from heights of 30 cm to 1 m, the protoplasts remained round and structurally intact, and chromosomes were not released (Fig. S1A). By increasing the dropping height to approximately 2.5 m, protoplast rupture occurred, releasing the cellular contents. Bright-field images showed large amounts of cell debris, while under fluorescence the entire cell lysate was stained with DAPI (Fig. S1B). To limit this background fluorescence, we stained the protoplasts with acriflavine (Raju, 1986) (Fig. S1C). We observed an average of 5.92 nuclei (range 2-12) per ungerminated conidiospore (Fig. 1A), while in the protoplasts derived from germinated spores, we observed an average of 2.26 (range 2-6; Fig. 1B). The fraction of binucleate protoplasts was higher than expected compared to the conidia from which they are derived, possibly resulting from the filtration steps eliminating larger cells with higher numbers of nuclei. Nevertheless, the numbers on nuclei that were observed in the protoplasts indicate that nocodazole indeed prevented nuclear division in the conidial germlings.

**Figure 1:**
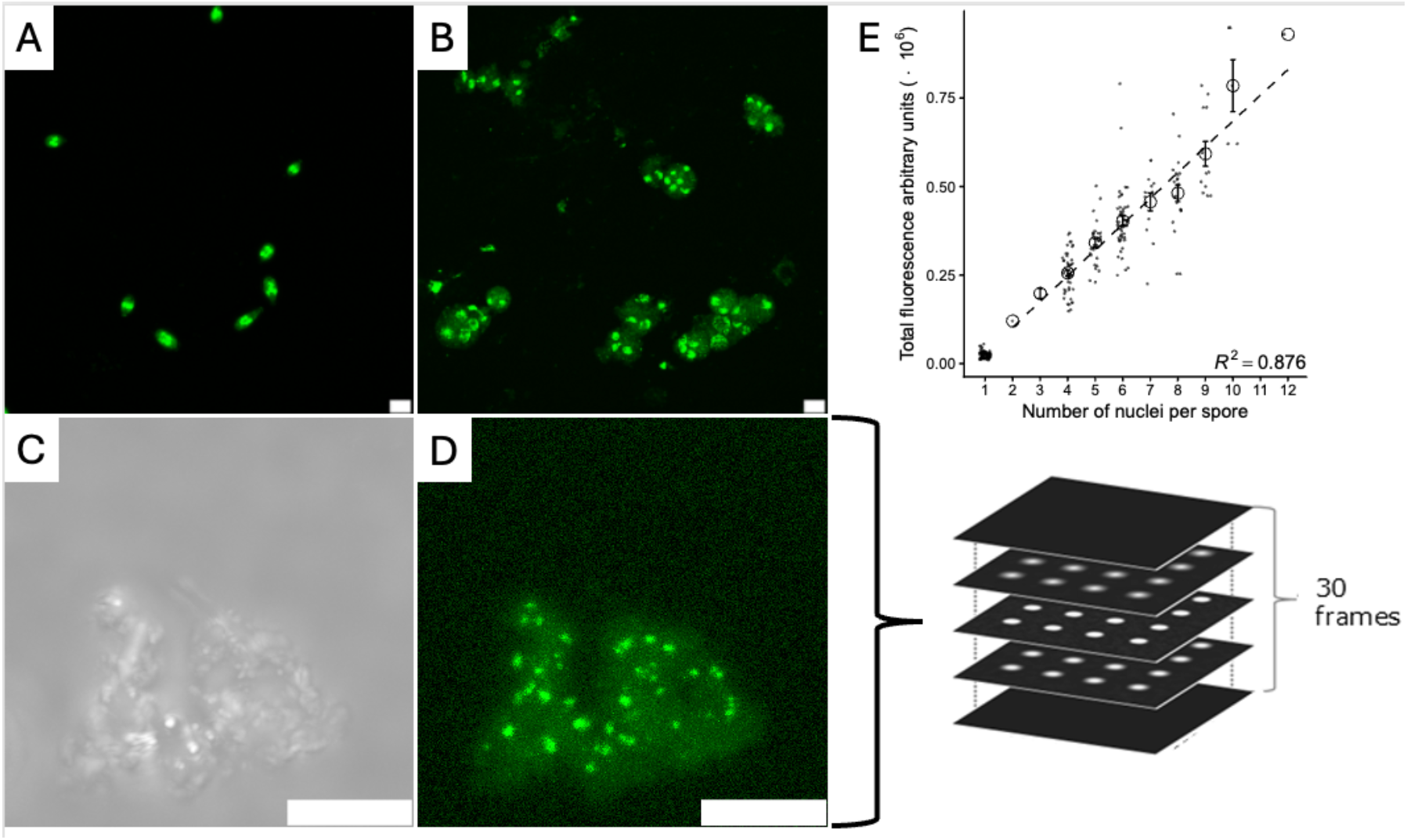
Nuclear and chromosomal distribution in *B. cinerea* indicate multiple chromosomal sets. Acriflavine stained conidiospores (A) and protoplasts derived from spore germlings (B) showing variable number of nuclei (green). (C) Bright-field images of the same protoplast as in (D). (D) Maximum projection image of acriflavine fluorescence of a single protoplast, with bright dots indicating chromosomes. (E) Relationship between the total fluorescence per spore and the nuclear number, with an increasing fluorescence based on number of nuclei. Dotted line indicates a linear model based only on conidiospores (i.e. spores with 2-12 nuclei), and not the microspores (mononucleate spores). Note: scale bars are 5 µm.

After physically dropping the protoplasts to rupture the plasma membrane (Fig. 1C+D), we observed more than 30 chromosomes per protoplast. As an example, Figure 1D shows a protoplast with 34 distinct fluorescence signals, which we infer to each represent a single chromosome (Movie S1). The *B. cinerea* genome contains 16 core chromosomes (>1.8 Mb) and 2 mini-chromosomes (<250 kb) (van Kan et al., 2017) and, although it is difficult to resolve precise chromosome counts, these results show that germlings of an individual conidiospore contain more than one haploid genome equivalent.

### Comparison of DNA content across spores with differing nuclear number

This new model suggests that multinucleate spores, like that of *B. cinerea*, have a constant DNA content regardless of their nuclear number. The significant variation in nuclear number we observed in the spores (Fig. 1A) allowed us to test this relationship. Across 180 spores, we found that the total nuclear acriflavine fluorescence of a spore was strongly correlated with the nuclear number (R^2^ = 0.88, *p* < 0.001; Fig. 1E; Movie S2), indicating increased DNA content with each subsequent nucleus.

The distribution of nuclear numbers in *B. cinerea* resulted in very few spores with low numbers of nuclei. To estimate the DNA content of a single-nucleus spore we leveraged microspores (spermatia), which are uninucleate non-germinating spores that are thought to be solely involved in sexual fertilization. To this end, we isolated a mutant which we have observed to display vastly increased microspore production and abolished conidiospore production, resulting from a premature stop codon in Bcin04g03490, a described developmental regulator. The average nuclear fluorescence of nuclei in these microspores was 2.42x10^4^ units, slightly lower than the 2.96x10^4^ units predicted by a linear model based on multinucleate spores alone (Fig. 1E). Despite potential differences in chromatin states and cell phases between these spore types, this level of fluorescence compares very well to that predicted by a linear relationship (Fig. 1E).

### Comparisons with *Neurospora crassa*

As this chromosome division model has recently been extended to the genetic model fungus *Neurospora crassa* (Tan et al., 2025), this provided an additional opportunity to test our results. The *N. crassa* spores are generally mononucleate or binucleate (Fig. 1A, Fig. 1B). Upon protoplasting germlings of conidia grown in the presence of nocodazole, we observed fewer chromosomes per protoplast than in *B. cinerea*, for example 14 in the *N. crassa* protoplast shown in Figure 2D. As there are 7 chromosomes in the *N. crassa* genome, this represents significantly more than one full set. A further strong indication that the fluorescent bodies in fact represent chromosomes comes from the observation that the chromosome in the top left has a “tail” visible in the confocal stacks (Movie S3). This feature has been reported previously for one of the chromosomes in *Botrytis* species, and it corresponds to the ribosomal DNA repeat (Shirane et al., 1988). Across 298 spores, we observed a similarly strong linear relationship between the nuclear number and the cumulative acriflavine fluorescence intensity (Fig. 1E) (R^2^ = 0.73, *p* < 0.001), indicating that nuclei in spores of *N. crassa* are likewise made up of repeating units.

**Figure 2:**
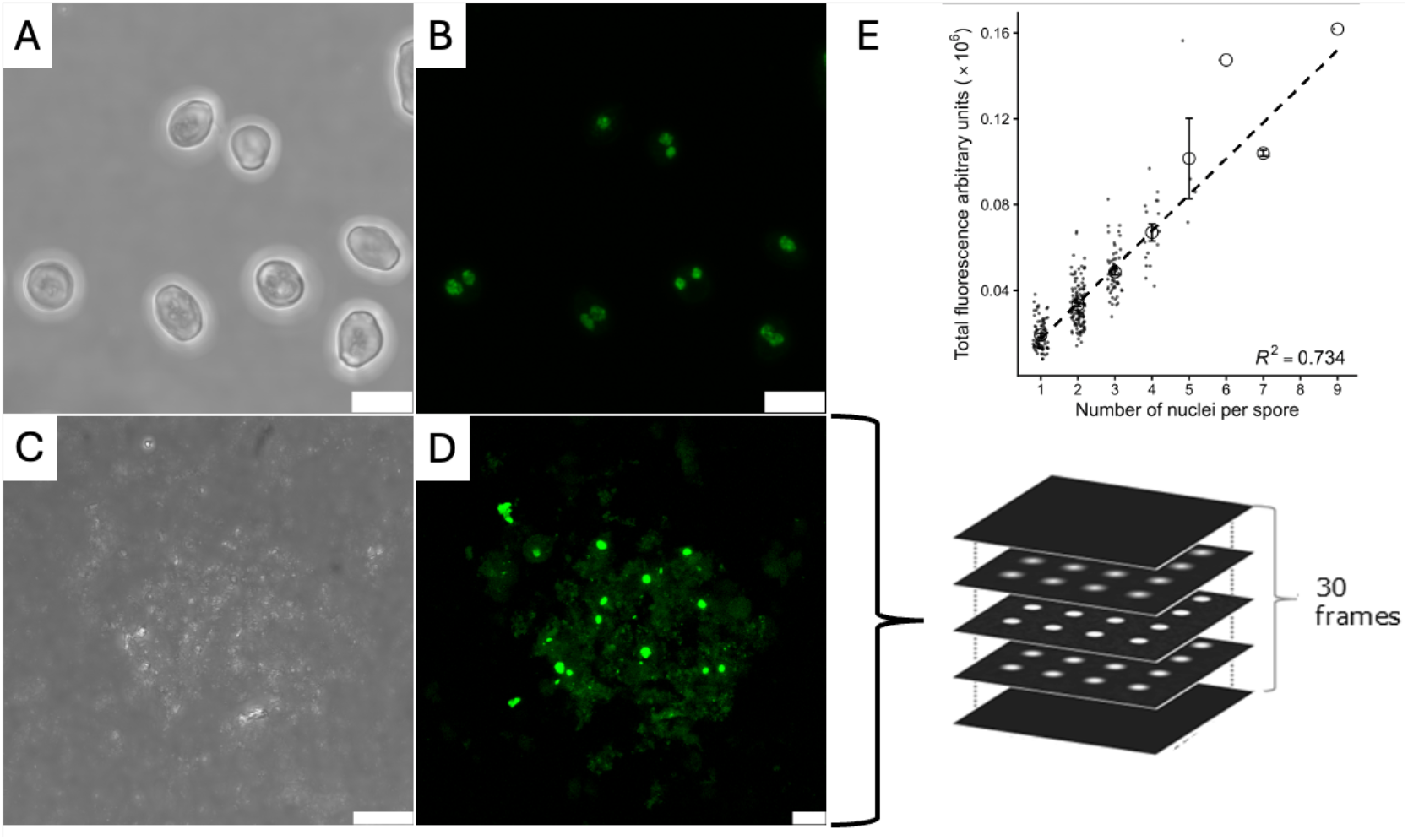
Nuclear and chromosomal distribution in *N. crassa* indicate multiple chromosome sets. Epifluorescence (A) and brightfield imaging (B) of acriflavine stained conidiospores showing variable number of nuclei (green). (C) Bright-field images of the same protoplast as in (D). (D) Maximum projection fluorescence of chromosomes from a single protoplast, with bright dots indicating chromosomes. (E) Relationship of the total fluorescence per spore and nuclear number, with an increasing fluorescence based on number of nuclei. Dotted line indicates a linear model Note: scale bars are 5 µm.

### UV-induced mutagenesis in *Botrytis cinerea*

The distributed genome model of Xu et al. (2025) makes an important prediction that mutations induced in a single spore will result in a colony where this variant is now fixed. Conversely, the traditional understanding of the multinucleate spores predicts that mutations will persist at an intermediate frequency, based on the number of nuclei inside a spore. To test this, we exposed conidiospores of *B. cinerea* to either 20 or 100 seconds of UV that resulted in 50 and 1% survival, respectively (Fig. 3A). To capture the genetic diversity of a single spore, we picked the entirety of individual germinated spores, harvested DNA from the resulting single-spore-derived colonies, and sequenced at an average coverage of 65X. To identify variants we kept only those found in a single sample, and not found in any of five replicate control samples that were not exposed to UV. We recovered an average of 18 and 66 *de novo* mutations in the 20 s and 100 s exposed spores, respectively (Fig. 3B; Fig. 3C). Of these, no mutation was fixed in a resulting colony, excepting a single mutations from one of the 100 s exposed spores (Fig. 3C). Much more frequently, we recovered intermediate frequency variants, with only a few greater than 0.5 frequency in a given colony.

**Figure 3:**
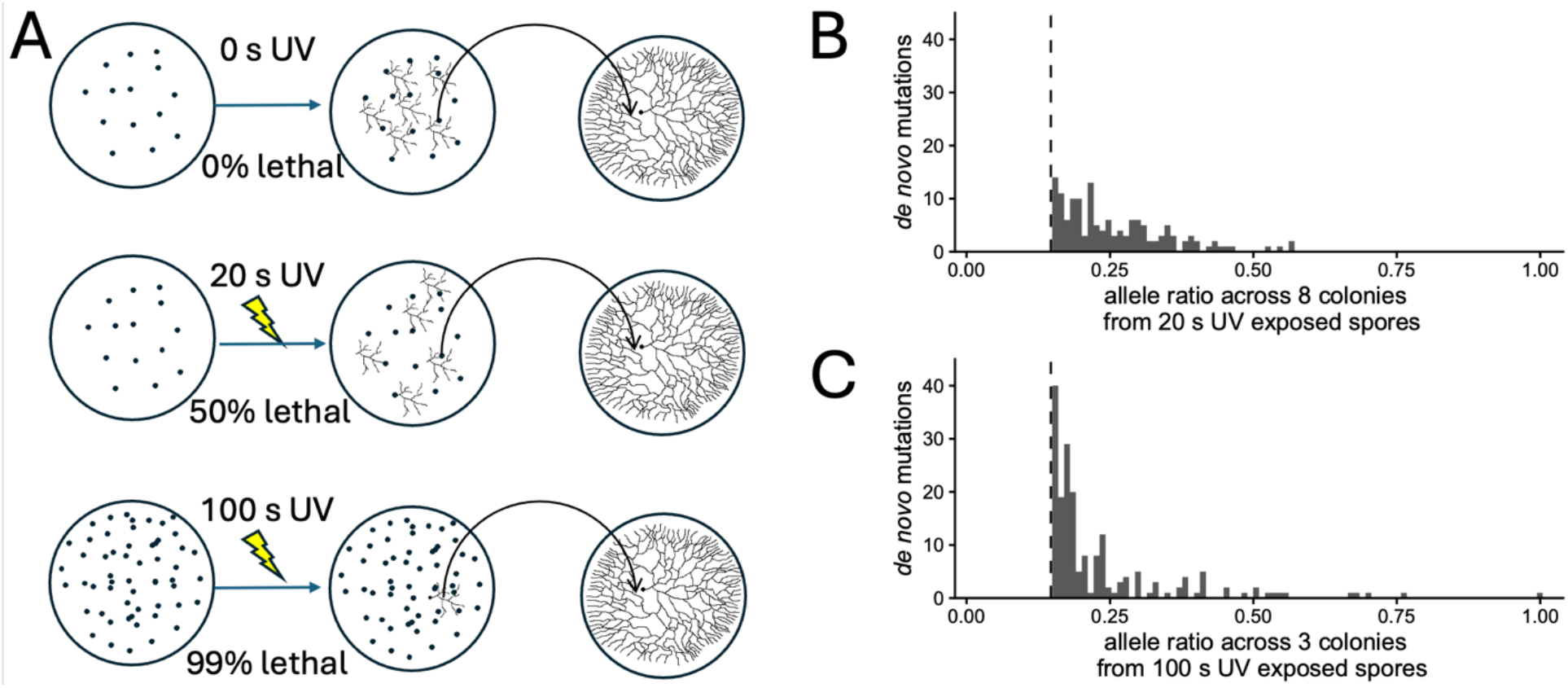
Whole genome sequencing of UV exposed spores does not recover evidence of fixed mutations. A) Experimental design to recover entire mycelia of colonies resulting from UV exposed spores. B) Allele ratios of *de novo* mutations across the 8 colonies from 20 seconds of UV exposure. Note that mutations with frequency < 0.15 (vertical dashed line) were not considered. C) as in B, but with 100 seconds UV exposure.

## Discussion

Our data indicate that, in two distantly related fungi, chromosome content of individual nuclei within multi-nucleate cells are fully haploid and do not comprise fractional complements of the genome. The most direct evidence is from chromosome counting in protoplasts. We counted multiple sets of the haploid number of chromosomes, in stark contrast to the microscopic evidence of a single haploid set shown for *S. sclerotiorum* spore-derived protoplast (Xu et al., 2025). To reconcile these findings, we note that in the previous images, the supposed protoplasts had a smooth surface with no remnants of cell walls and a complete absence of ribosomal aggregates or organelle structures. In contrast, our images display clear evidence of cell component remnants and other irregular structures, being multiple microns thick. Thus, we do not disagree with the count itself, but rather the unit of observations. We interpret the data of Xu et al. (2025) to represent free nuclei released during the protoplasting procedure (Tallman & Reeck, 1980). This interpretation is consistent with the difficulty in preparing burst protoplasts for microscopy, as we required heights greater than 3 m, while the authors of Xu et al. (2025) communicated that 0.3 m was sufficient in their experiments.

In *B. cinerea* protoplasts, we observed a significantly larger number of chromosomes than a single set, roughly equivalent to two haploid genome copies. A microscopic observation of chromosomes in *Botrytis cinerea* and five other *Botrytis* species was documented in the 1980’s by Shirane et al. (Shirane et al., 1988, 1989). From hyphal nuclei, these authors generated a full microscopic metaphase karyotype of all six species, five of which contained 16 chromosomes while one species, *B. allii*, contained 32 (Shirane et al., 1989). The latter species was later shown to be an allodiploid hybrid species between *B. byssoidea* and *B. aclada* (Nielsen & Yohalem, 2001). As this work bypasses the need for protoplasting and nocodazole treatment, it provides additional support that, at least in hyphae, nuclei contain full chromosome sets.

The original motivation for the work in *B. cinerea* and *S. sclerotiorum* was said to be the observation of morphologically homogenous mutants in *S. sclerotiorum* in a UV screen (Xu et al., 2022), which the authors suggest can be explained by the sorting of chromosomes between nuclei (Xu et al., 2025) making spores effectively haploid. Testing this hypothesis, we mutagenized *B. cinerea* spores with UV light. We did not find strong evidence of fixed mutations in the resulting colonies. We were able to generate significant genetic variation although identifying high-confidence low-frequency variants is difficult, the amount of induced mutations was proportional to the amount of UV exposure, indicating our results are not solely bioinformatic artefacts. Our results are consistent with multiple, redundant, haploid nuclei inside a single spore, with mutations occurring in a single nucleus and only reaching intermediate frequencies. A more rigorous test of this hypothesis would require whole genome amplification of single nuclei, followed by genome sequencing. This method has been used in arbuscular mycorrhizal fungi, the extreme of multinucleate fungi, where the results clearly support a full set of chromosomes in each nucleus (Chen et al., 2018; van Creij et al., 2023). If such amplifications are attempted, it is important to be aware of the peculiar uneven coverage of fungal whole genome amplifications when drawing inferences. Often, amplification of single nuclei only covers 30-50% of the genome, and the extensive missing data requires careful examination (Auxier & Bazzicalupo, 2019). This uneven amplification coverage may explain the PCR results shown for both *B. cinerea* and *S. sclerotiorum*, where not all chromosomes were PCR-positive (Xu et al., 2025). An important technical detail to note here that in the work of Xu et al. (2025) the two markers for each chromosome, for the left and right arms, the primers were designed to produce products of the same size and then run in multiplex in the same reaction tube. Thus, this assay does not demonstrate the presence of entire chromosomes, as it cannot be ascertained if one or both arms were amplified. If the entire genome is not amplified from a single nucleus, there will be insufficient target for subsequent PCR amplifications.

Together, our evidence is fully consistent with previous assumptions about the genetic makeup of multinucleate fungi. Outside of these experiments, perhaps the most commonly encountered evidence for multiple haploid genomes in spores is observed when making transformants in *N. crassa*. In this species, it is routine to directly electroporate conidia and, like the steps that Beadle and Tatum used, the resulting heterokaryotic primary mutants must be purified to a homokaryon through a sexual cycle (Chakraborty et al., 1991; Colot et al., 2006). This is a tedious undertaking, and often the bottleneck in larger molecular biology projects in this species. While a distributed chromosome model is intriguing, it must account for all available evidence, not only selected experiments.

## Supplemental Figures

**Figure S1:**
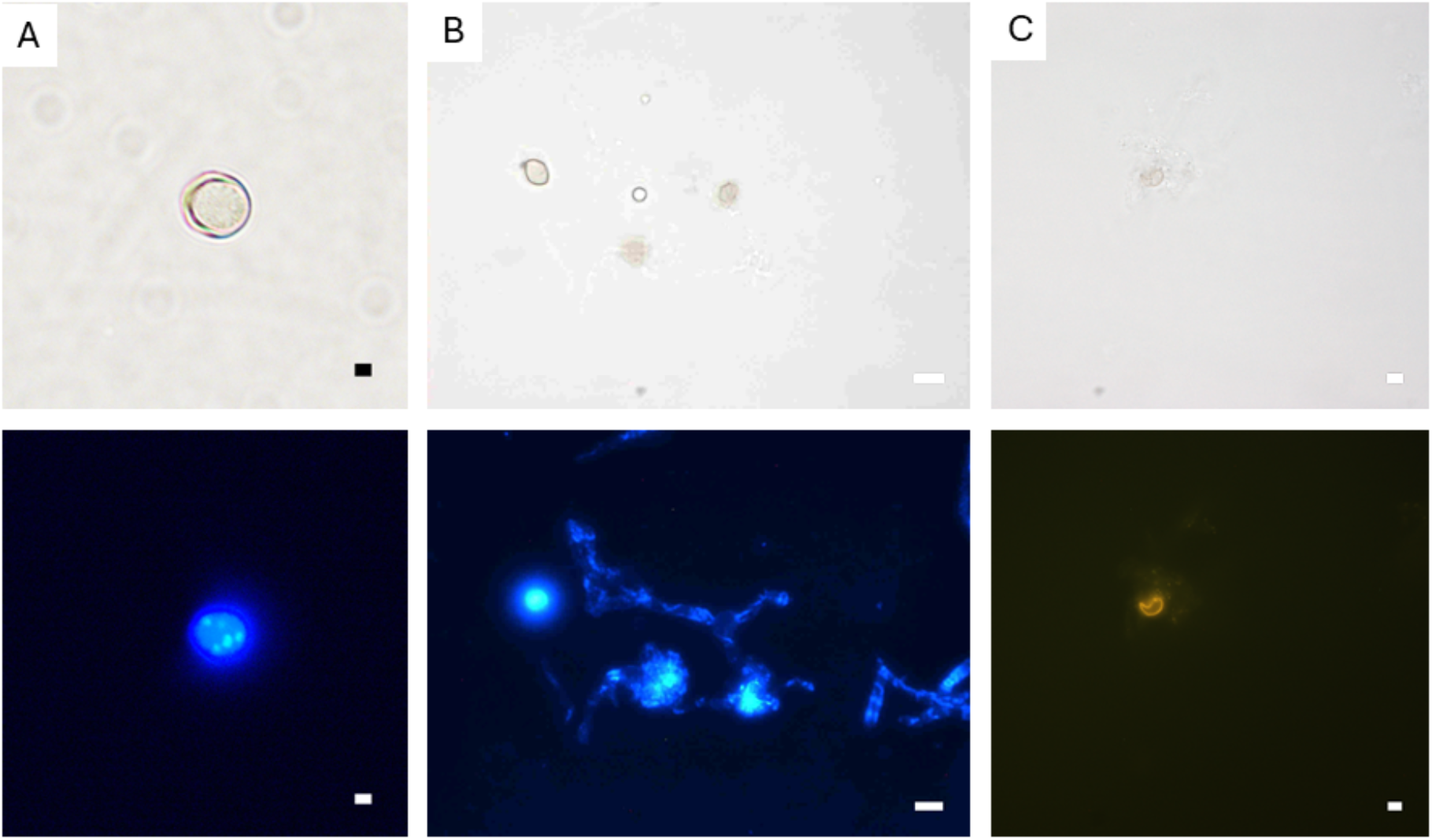
Dropping and staining for observation of chromosomes. A) Dropping of protoplasts from 1 meter high results in intact protoplasts. Top panel is bright field microscopy with fluorescence microcopy using DAPI below. B) Dropping from a height of 2 meters can release the nuclei from the protoplasts, but DAPI signal (bottom) is obscured by cytoplasmic fluorescence. C) Dropping from 2.5 meters high released the chromosomes from protoplast, and acriflavine shows increased specificity in staining (bottom).

**Figure S2:**
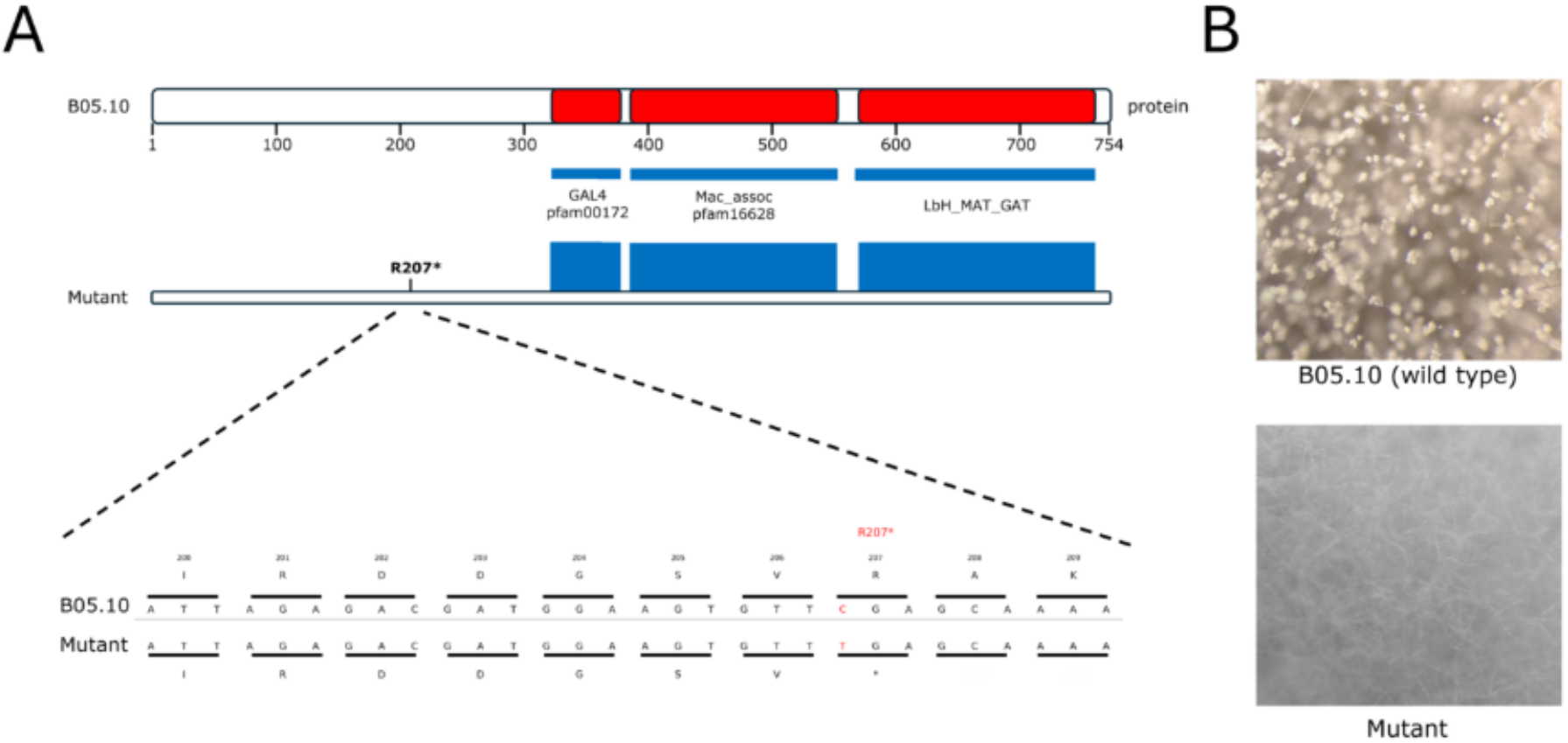
Mutation in Bcin04g03490 resulted in abolished macroconidia. A) Protein domains of Bcin04g03490. The mutation R207* created a premature stop codon, prior to any of the predicted protein domains. B) B05.10 (wild type) exhibited normal conidiophore formation, while no conidiophores are observed in the mycelium from the mutant.

**Supplemental Movie S1**: Combination of all the acquired 37 images of chromosomes from dropped protoplast from *B. cinerea* stained with acriflavine. The movie progresses from the lowest Z-position with a speed of 1 slice per second to the highest Z-position.

**Supplemental Movie S2**: Combination of all the acquired 78 images of conidiospores with different numbers of nuclei from *B. cinerea* stained with acriflavine. The movie started from the lowest Z-position with a speed of 3 slices per second to the highest Z-position.

**Supplemental Movie S3**: Combination of all the acquired 30 images of chromosomes from dropped protoplast from *N. crassa* stained with acriflavine. The movie progresses from the lowest Z-position with a speed of 1 slice per second to the highest Z-position.

## Methods

### Strains and culture conditions

The *B. cinerea* strains used in this study were B05.10 and mutant SAC24 (abolished conidiospores production). Strains were cultured on Malt Extract Agar (MEA, OXOID) plates at 20 °C in the dark. To produce conidiospores, plates were exposed to black light as previously described (Zhang et al., 2016). For the production of microspores, plates were first inoculated at 20 °C in the dark for 1 month and then transferred to 0 °C incubator darkness for another month incubation before the microspores were harvested.

For *N. crassa* observations, a standard reference strain, FGSC2489, was cultivated under standard conditions. For conidia production, strains were cultured at 25 °C with 16 hours of light.

### Protoplasting of both *B. cinerea* and *N. crassa*

To prepare protoplasts 1x10^8^ asexual conidiospores were inoculated in 100 ml of HA medium (4 g/L Glucose, 4 g/L Yeast extract, 10 g/L Malt extract, pH 5.5) with 100 µL Nocodazole. Spores were germinated by overnight incubation with shaking in a 250 mL Erlenmeyer flask (ca. 18 h; 20 °C; 180 rpm). The liquid was transferred into two 50 mL tubes and centrifuged at 1,000 g for 8 min with a swinging bucket rotor. The supernatant was discarded supernatant, resulting in mycelium pellets of ∼3-4 g fresh weight, which were washed twice with 40 mL of 0.6 M washing buffer (0.6 M KCL, 0.1 M Na_2_HPO_4_, 0. 1M NaH_2_PO_4,_ pH 5.8), followed by centrifugation for 5 min at 1000 g. The pellet was resuspended in 20 mL Protoplasting Enzyme Mix (0.6 M KCl, 0.1 M Na_2_HPO_4_, 0.1M NaH_2_PO_4_, 10 mg/ml Vinotaste pro), and incubated on a Rocky 3D shaker (VWR International) at intensity 4.5-5 for 60-90 min at 28 °C. Protoplasts were filtered through sterile nylon cloth (30 µm) into 10 ml ice-cold TMS buffer (1M Sorbitol, 10 mM MOPS, pH = 6.3). This was then pelleted by centrifugation for 4 min at 4 °C, 400 g, and the supernatant was removed.

### Chromosome visualization

To fix protoplasts, 800 µL of 3:1 methanol:acetic acid solution was added to resuspend the pelleted protoplast and transferred to 2 mL tubes. Tubes were placed on a horizontal roller for 1 hour. This mixture was centrifuged at 400 g for 4 minutes, the supernatant was discarded and the protoplasts were hydrolyzed in 4 N HCl for 100 min at 30 ºC. This was then centrifuged at 400 g for 4 minutes and resuspended in 1 mL of distilled water. These protoplasts were then centrifuged at 400 g for 4 minutes and resuspended in 90% ethanol with 1 µL of acriflavine stock (200 mg/mL) and incubated for 60 mins at 30 ºC. This was washed three times at 30º C for 5 minutes with 1mL concentrated HCl-70% ethanol mixture to remove non-covalently bound stain. Finally, the stained protoplasts were centrifuged at 400 g for 4 minutes resuspended in water. To visualize chromosomes, 200 µL of the fixed protoplasts were dropped onto glass slides in a large petri dish with a height 3 m, and air dried for 20 minutes. 20 µL distilled water was added on the sample and a #1.5 coverslip (BRAND^®^) was gently placed on the top.

Initially, the stained dropped protoplast were examined with an epifluorescence microscope (Nikon 80i) employing an HBO 100 mercury lamp (Fig. S1). For DAPI staining (Fig. S1 A+B), the excitation filter/dichronic beam-splitter/barrier-filter combination was chosen for excitation at 350 nm and emission at 420 nm (DAPI filter cube; EX 340-380 DM 400 B A 420. For acriflavine staining (Fig. S1C), the excitation filter/dichroic beam-splitter/barrier-filter combination was chosen for excitation at 460 nm and emission at 510 nm (GFP-L filter cube; EX 460-500 DM 505 B A 510). Observations are made at a magnification of 500× or higher.

Confocal observations (Stellaris 5 Confocal LSM Leica) were made with a 100X oil-immersion objective with excitation at 450 nm and emission 480–600 nm. Z-stacks were acquired in 0.33 µm steps, with a scanning diameter of 1 Airy Unit. For single protoplasts, a maximal projection image of 37 Z-stack layers was generated with Leica Application Suite X, version: 3.10.1.29575. The numbers of chromosomes were counted by ImageJ (version 1.51f), using the following parameters: 1. Image Type: 8-bit; 2. Process filters: Gaussian Blur sigma =1; 3. Image Adjust Threshold: Otsu 18,255 darkbackground; 4. Analyze Particles: Size: 3-indefinite, Circularity: 0.2 - 1.0, Show: Outlines, Display Result, Add to Manager.

### Acriflavine staining to quantify the total amount of DNA in spores from *B. cinerea* and *N. crassa*

In *Botrytis cinerea*, 2 different types of spores were used in this experiment: conidiospores and microspores. Conidiospores are harvested from the plate following the protocol of You et al. (2024), modified by a centrifuge speed of 4000 g for 10 minutes in each step. Microspores and conidiospores were diluted to 1x10^7^ /mL, and 100 µL from each spore suspension was mixed in a microcentrifuge tube. Spores were stained with acriflavine as described above. The observation for both *B. cinerea* and *N. crassa* under confocal microscopy was the same as described above.

To calculate the fluorescence signal intensity per spore in both *N. crassa* and *B. cinerea*, we manually selected nuclei using Fiji version 2.17.0 with following steps: 1. Bio-Formats Importer: Hyperstack, color mode: Grayscale; 2. Z project: Projection type = Sum Slices; 3. Freehand selections: select nuclei within the spore; 4. Analyze: Measure. Integrated Density per spore = Area x Mean Intensity.

### Determination of UV induced mutation frequency

To test the effect of UV, we plated conidiospores of *B. cinerea* at a low density on MEA and immediately exposed these plates to either 0, 20, or 100 seconds of UV light (1620 µW/cm^2^). Following 2 days of incubation at 20 ºC under darkness, we identified isolated growing colonies originating from a single spore with a Leica DMi1 inverted microscope, and transferred the entire colony to a new plate. This plate was incubated, and all mycelia was harvested from the plate, to capture the genetic variation of the colony. DNA was extracted as previously described (Rodenburg et al., 2018). This DNA was prepared for sequencing using the HackFlex protocol, adapted for 96-well plates (Gaio et al., 2022; Reyes Márquez et al., 2025). The resulting library was sequenced on a Illumina XXX, producing 150 bp paired-end reads.

These reads were aligned to the reference genome (ASM83294v1) (Amselem et al., 2011; van Kan et al., 2017) with bwa-mem2 v2.2.1, and variants were predicted with GATK v4.2.6.1 (Auwera & O’Connor, 2020; Vasimuddin et al., 2019). Variants were filtered with recommended hard filtering, and high-confidence *de novo* mutations were identified with vcfR (Knaus & Grünwald, 2017) based on having a minimum read depth of the mean depth of the sample, and being unique to a given sample (i.e. no alternate reads for a given position in other samples).

### Data and Code Availability

Code for analysis of nuclear fluorescence and *de novo* mutations can be found at https://github.com/BenAuxier/chromosome_sets. Raw DNA sequencing data for *B. cinerea* mutations can be found at PRJEB110055

## Acknowledgements

We thank Francisca Reyes Marquez for assistance with DNA sequencing. We also thank Jeff Rollins for critical feedback on the manuscript.

## Notes

### Competing Interest Statement

The authors have declared no competing interest.

